# A role of cryptochrome for magnetic field-dependent improvement of sleep quality, lifespan and motor function in *Drosophila*

**DOI:** 10.1101/2021.08.02.454845

**Authors:** Haruhisa Kawasaki, Hideyuki Okano, Hiromi Ishiwatari, Tetsuo Kishi, Norio Ishida

## Abstract

Understanding the molecular genetic basis of animal magnet reception has been one of the big challenges in biology. Recently it was discovered that the magnetic sense of *Drosophila melanogaster* is mediated by the ultraviolet (UV)-A/blue light photoreceptor *cryptochrome* (*Cry*).

Here, using the fruit fly as a magnet receptive model organism, we show that exposure to a specific AC magnetic field during night time affected the health of the fly. AC magnetic field exposure showed lifespan extension under starvation, sleep improvement and prevention of decreased motor function. In contrast, all the health improvement effect was not observed in *cryptochrome* mutant flies (*cry*^*b*^). We showed that AC magnetic field exposure prevented motor dysfunction in Gaucher’s disease model *Drosophila*. The data suggests that magnetic field-dependent improvement of sleep quality, lifespan and motor function is mediated through a *cry*-dependent pathway in animals.

## Introduction

All living things have been constantly exposed to the natural magnetic field. The exact effects evoked are not well understood, but exposure appears likely to cause biological effects. Recently, several organisms, such as cockroaches^1^, flies^2-4^, butterflies^5^, birds^6,7^ and humans^8^ detect and exploit magnetic field through one of the biological clock genes, *cryptochrome* (*cry*). Nerve-specific *cry* expression in fruit fly, *Drosophila melanogaster* increased magnetic field sensitivity^9^. A specific geographic magnetic field is imprinted on *Drosophila* and is inherited to its progeny^10^. The *cry* gene plays a critical role in magnetic detection among diverse species. Qin et al reported that CRY protein forms complexes with MagR protein in the retina of migratory birds suggesting they could sense the Earth’s magnetic field^2^.

Organisms can not only detect magnetic field, but also get health improvements such as blood pressure and bone mineral density by exposure to magnetic field^11-13^.

Here, we report the results of the evaluation for alternating current (AC) magnetic field effect on health improvement by using *Drosophila melanogaster*. AC magnetic field exposure showed the lifespan extension under starvation, sleep improvement and prevention of decreased motor function. Furthermore, motor dysfunction in neurodegenerative disease model flies was prevented by AC magnetic field exposure.

## Materials and methods

### Flies

We used Oregon-R as the wild-type strain and *cry*^*b*^ as a *cry* mutant. CG31414 is a model fly for Gaucher disease^14,15^. These flies were raised under a 12-h light/12-h dark cycle at 25°C.

Standard culture medium consisted of 8% corn meal, 5% glucose, 5% dry yeast extract, 0.64% agar were boiled and supplemented with 0.5% propionic acid and 0.5% butyl p-hydroxybenzoate. For the starvation assay, flies are kept in 1% agar medium.

### Magnetic field exposure

In this research, 50 Hz magnetic field exposure was conducted using an AC magnetic field exposure device (Soken MS, Toride, Ibaraki, Japan), which has two separate magnetic coils inside the device^16,17^. The value of peak magnetic flux density *B*_max_ is 180 mT and *B*_rms_ is 127 mT on the surface of the magnetic field exposure device above the center of the coils.

### Assays of sleep behavior

Sleep behavior was recorded as described previously^14^. Male flies were placed in glass tubes (inner diameter, 3 mm) containing 5% sucrose and 2% agarose and exposed to the magnetic field. The tubes were then transferred to a *Drosophila* Activity Monitoring (DAM) system (TriKinetics, Waltham, MA, USA). Locomotor activity was measured in one-minute bins and sleep was traditionally defined as ≥ five minutes of consolidated inactivity^18^. Sleep analysis software provided by M. Shimoda (National Institute of Agrobiological Science) was used to analyze the *Drosophila* locomotor activity data and sleep data. All experiments were tested by using male flies.

### Climbing assay

The climbing assay was conducted as described previously^14,19,20^. In brief, groups of 10 male flies per vial (2.5 cm diameter) were gently tapped to the bottom and the number of flies crossing a line at 6 cm height within a time period of 10 s.

## Results

### Determination of the optimal magnetic field strength

Validation of the optimal magnetic field strength is required to evaluate magnetic field therapy for health. Then we estimated magnet field decay with distance from AC (Alternating Current) magnetic field generator (Figure 1a). In this way, AC magnetic field intensity is attenuated with distance from the field generator gradually. In turn, we investigated the intensity of AC magnetic field which prolongs *Drosophila* lifespan under the starvation condition (Figure 1b). Though the lifespan of *Drosophila* is usually 2 months, it will be reduced to 3 days under the starvation condition while keeping humidity. For this assay, we altered the distance between the magnetic field generator and a vial containing *Drosophila* in a stepwise fashion and found that wild type *Drosophila* was significantly extended its lifespan at magnetic field strength of 0.5 – 0.7mT. In contrast, we did not observe similar extension in experiments by using *cry* mutant *Drosophila*.

**Figure 1.**
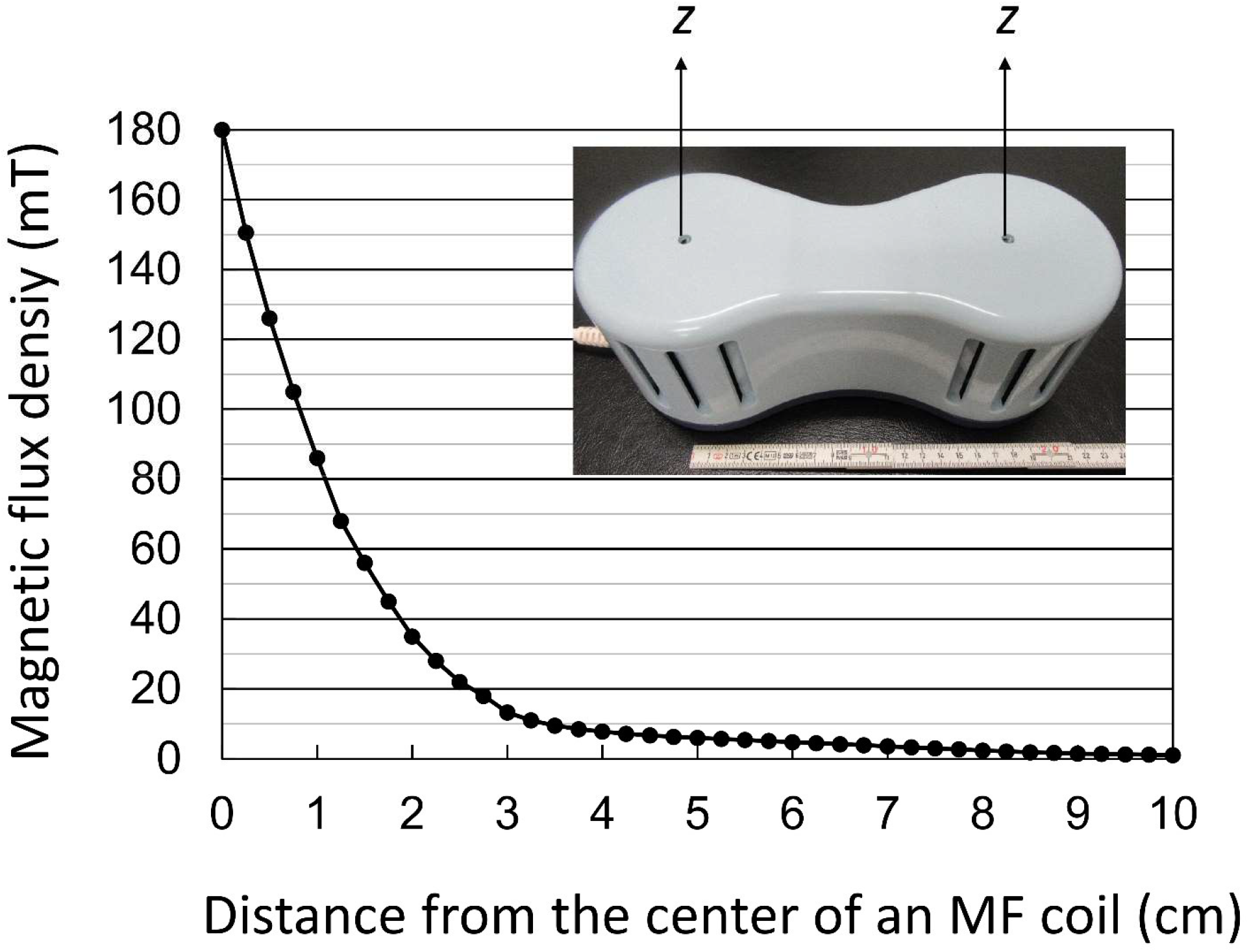
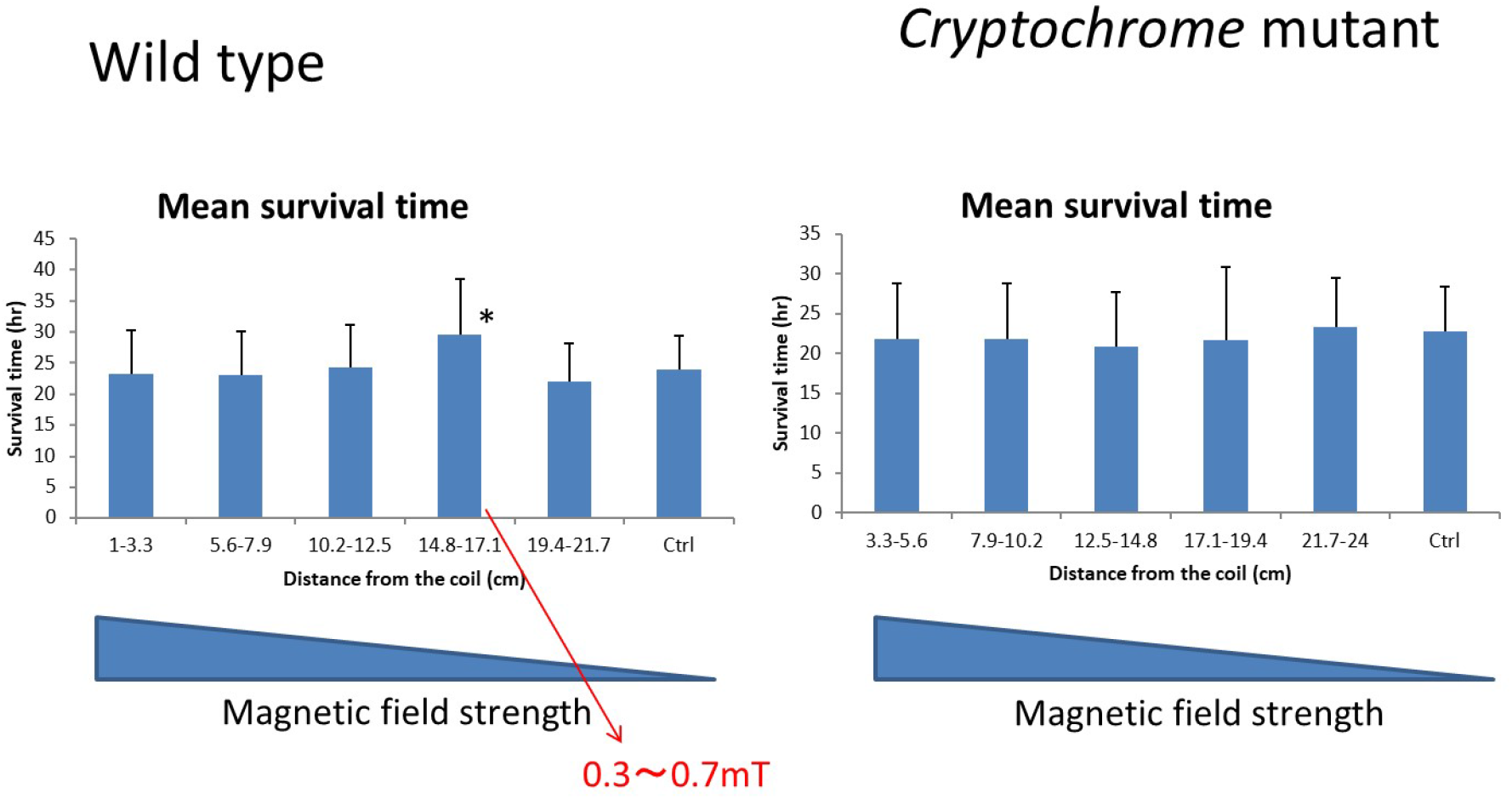
Determination of the optimal magnetic field strength. a. Relationship between magnetic flux density and distance from the center of a magnetic field (MF) coil. The MF strength decays with distance from the coil ^16,17^. b. Examination of optimal conditions for health promotion. Optimized distance for health promotion was determined by altering the distance from the coil. In comparison to the control group, flies at the distance of 14.8-17.1 cm from the center of an MF coil (that is, the MF strength of 0.3-0.7 mT) significantly extended lifespan in wild type fly.

### Magnetic field exposure increased motor function in wild type and neurodegenerative disease model flies

Gaucher disease is a genetic disease with neurodegeneration and *Gba1b* is the orthologue of Gaucher disease causative gene^14,15^ in *Drosophila. Gba1* mutation flies (CG31414) indicate some symptoms similar to humans, such as sleep disorder, motor dysfunction, and decreased lifespan^14^. We put Gaucher’s disease mutant CG31414 and wild type flies in the starved condition and exposed AC magnetic field (0.3-0.7mT strength) to determine the motor function by using climbing assay after 16 and 23hr magnetic exposure (Figure 2a). Flies under starved condition declined motor function, but AC magnetic field treatment increased motor function in both wild type and neurodegenerative flies (Figure 2b).

**Figure 2.**
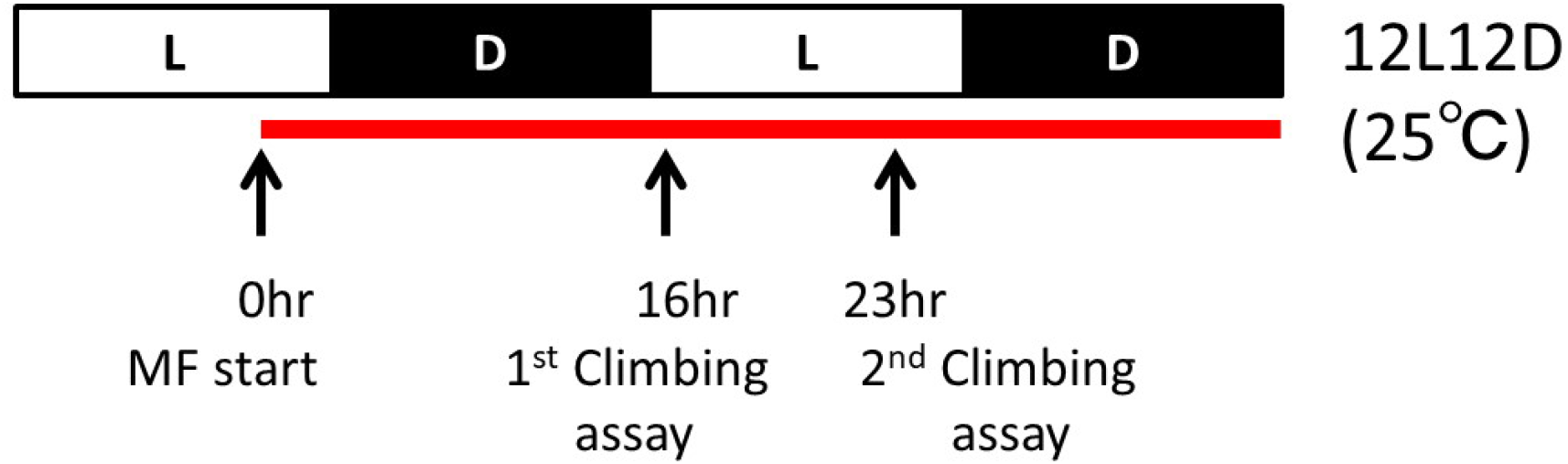
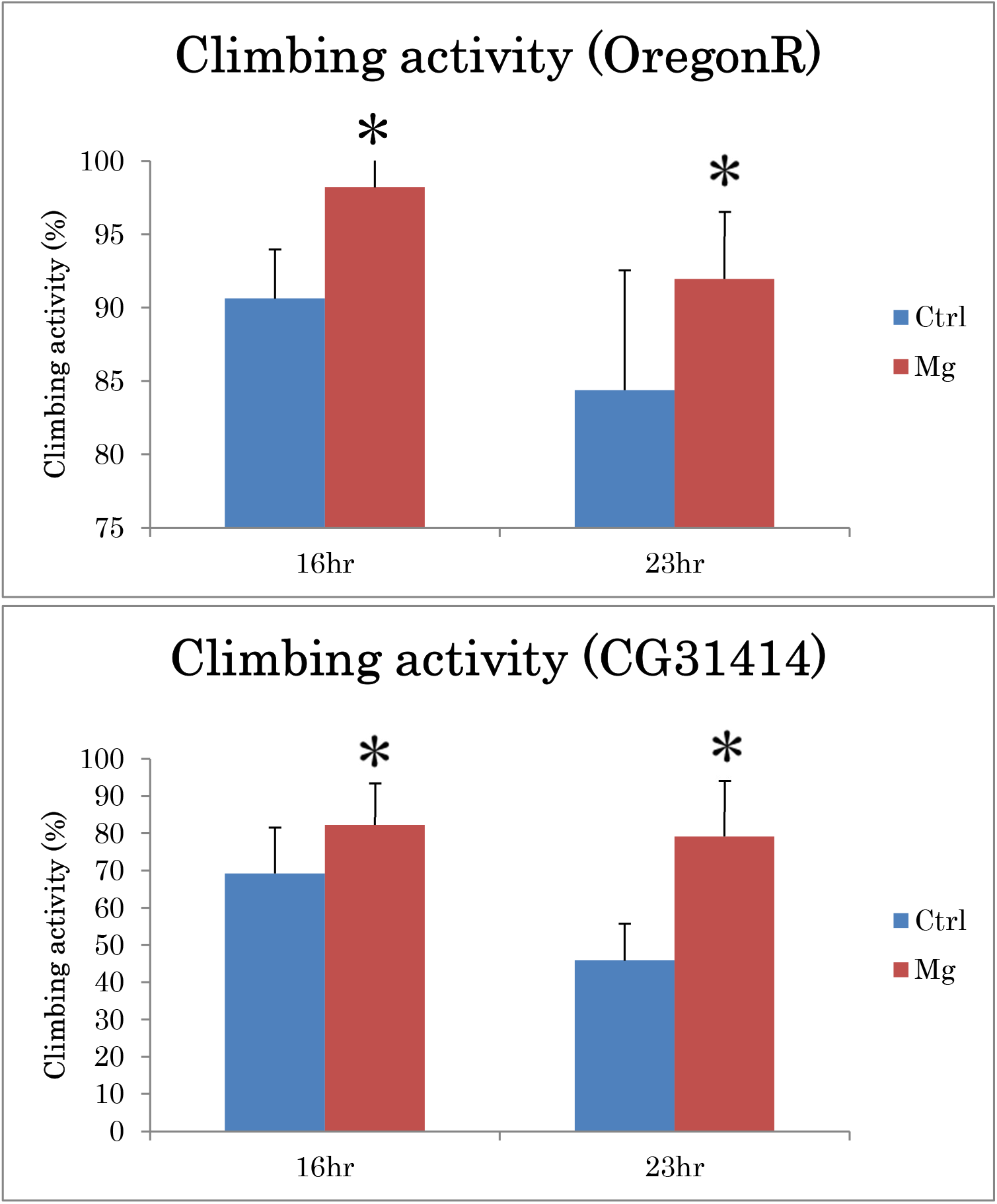
Magnetic field exposure increased motor function in wild type and neurodegenerative disease model flies. a. Experimental scheme for climbing assay. Flies 1-2 days after eclosion were put in starved condition and exposed AC magnetic field and determined climbing activity at 16 and 23 hr after magnetic field exposure. b. Climbing activity for wild type (Oregon R) and neurodegenerative disease model flies. CG31414 is Gaucher’s disease model. We compared climbing activity for control and magnetic field exposed flies. Statistical data by Student’s t-test are expressed as mean ± SD. * represents significant differences (p < 0.05).

### Sleep improvement by AC magnetic field exposure at night

Next, we evaluated the effects of AC magnetic field exposure for sleep in *Drosophila*. As shown in Figure 3a, we attempted sleep analysis by DAM system after AC magnetic field exposure at daytime or nighttime. These data indicated that wild type *Drosophila* flies exposed AC magnetic field at night time decreased sleep bout number and increased sleep bout length, in contrast, day time AC magnetic field exposure did not change sleep bout number and sleep bout length. In other words, the data indicates that night time AC magnetic field prevents sleep fragmentation and improves sleep quality. Furthermore, we did not observe such sleep improvements in *cry* mutant flies (Figure 3c).

**Figure 3.**
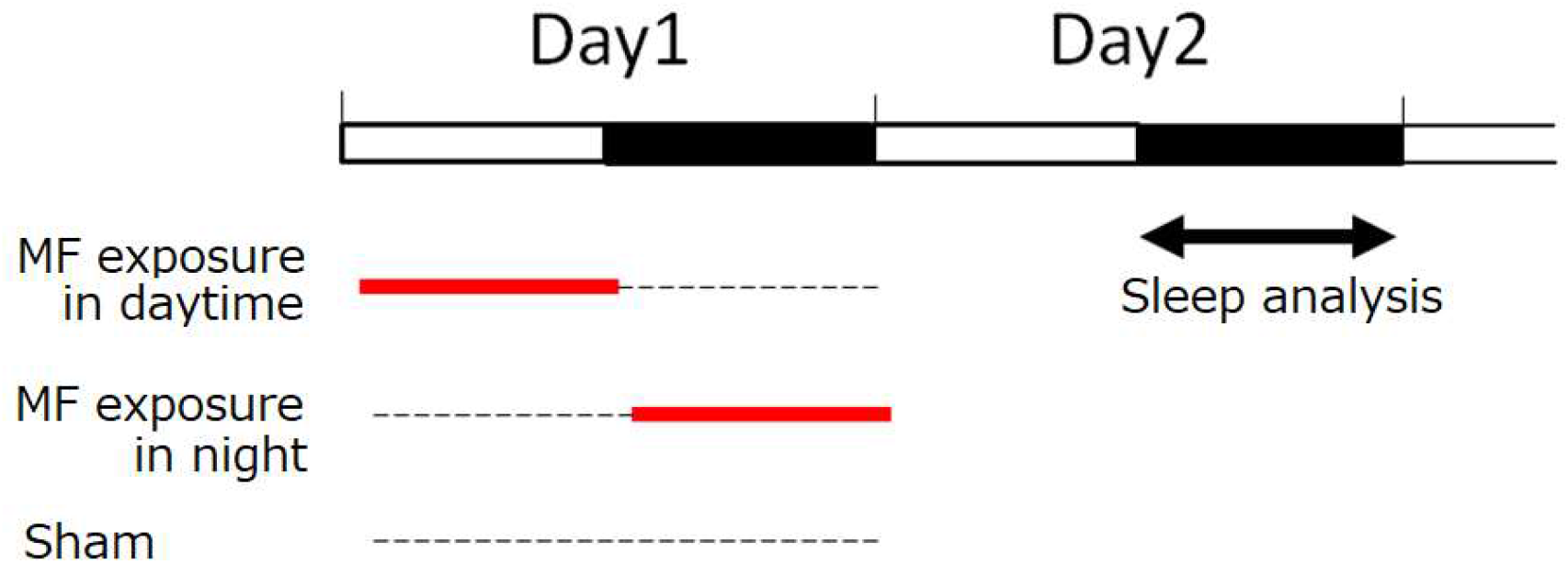
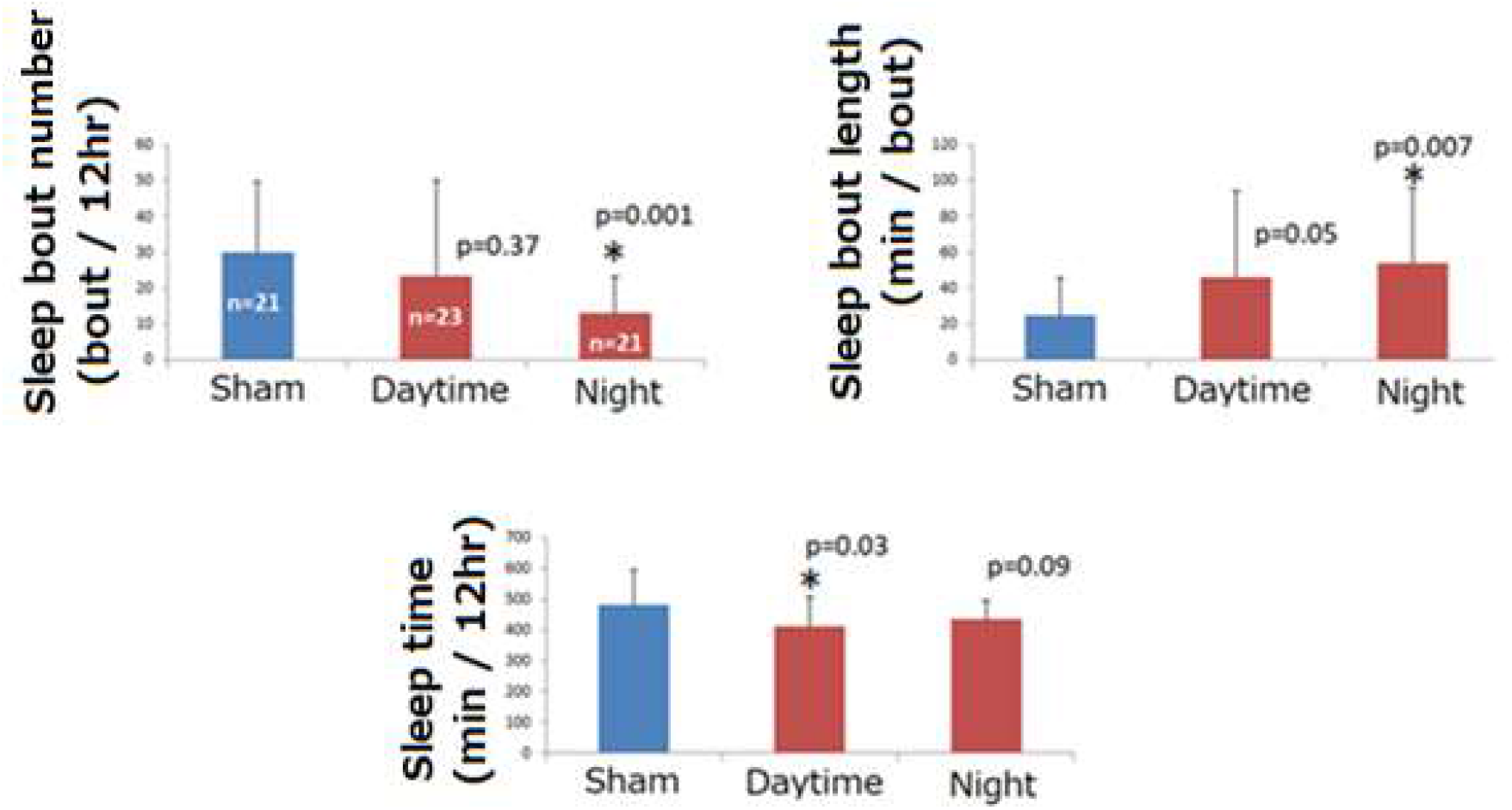
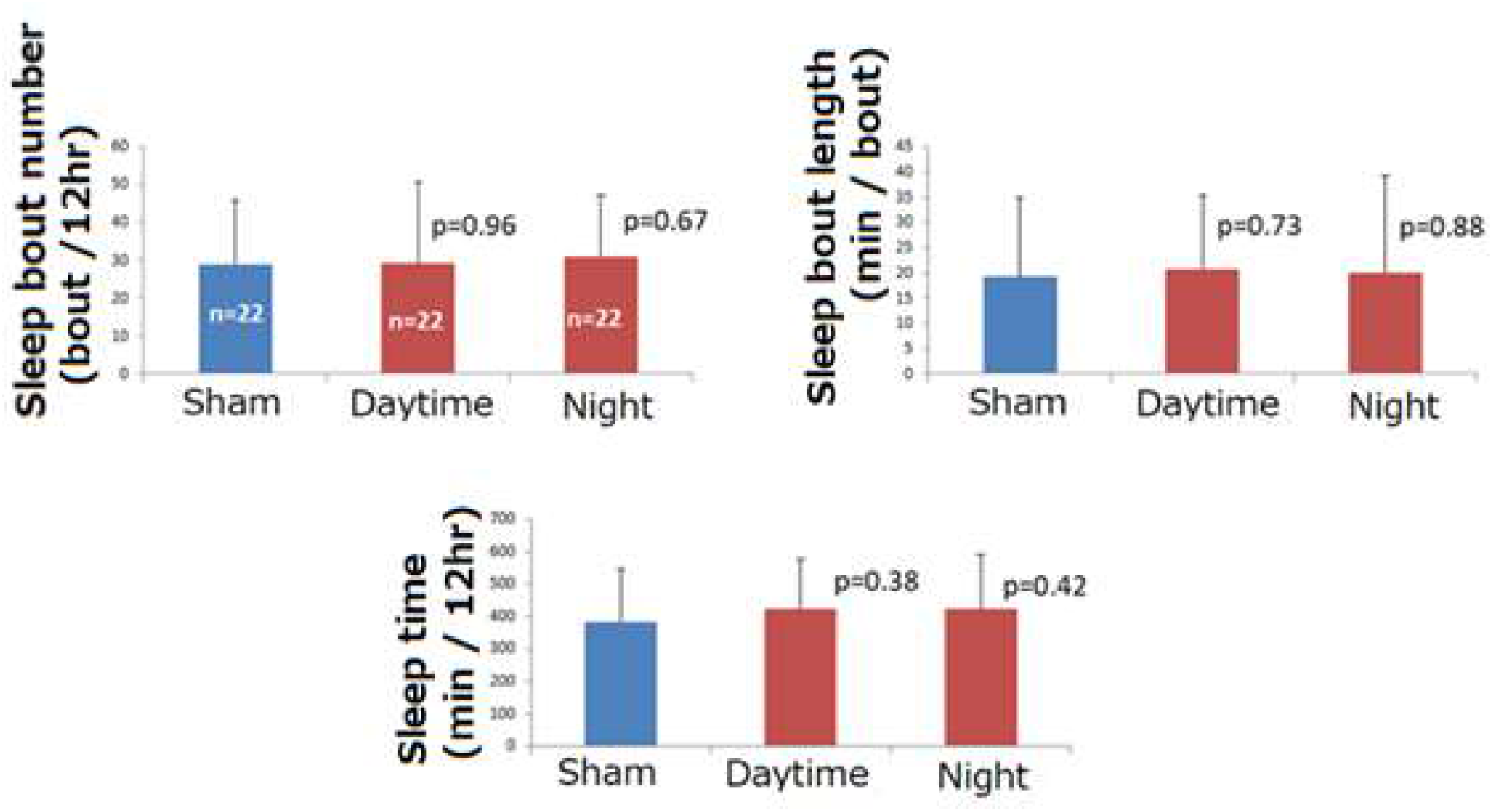
Sleep improvement by AC magnetic field exposure. a. Experimental scheme for sleep analysis after AC magnet field exposure (Shown in red). The duration of sleep analysis was shown in black arrow. b. Sleep analysis data for wild type flies. c. Sleep analysis data for *Cry* mutant (*cry*^*b*^) flies.

### Repeated magnetic field exposure increased motor function through *cryptochrome*

Our data revealed that nocturnal AC magnetic field treatment improves sleep quality, then we investigated whether this treatment affects other health promotion. We evaluated motor function by climbing assay for flies under starvation after repeated nocturnal AC magnetic field treatment three times. 11-15 days after eclosion male flies received night time AC magnetic field exposure repeated three times then transferred to nutrient-starved condition (Figure 4a). The climbing assay was performed at 15 and 23 hr after they were starved.

**Figure 4.**
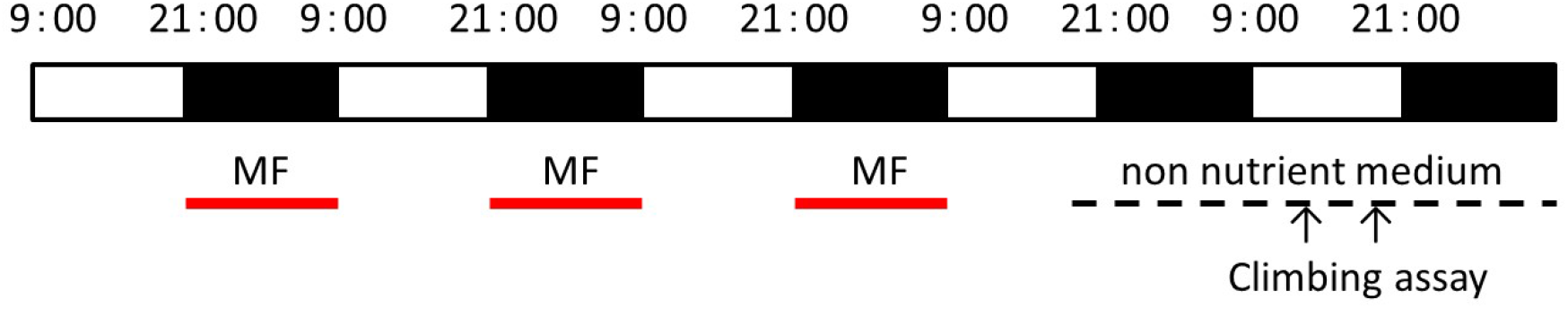
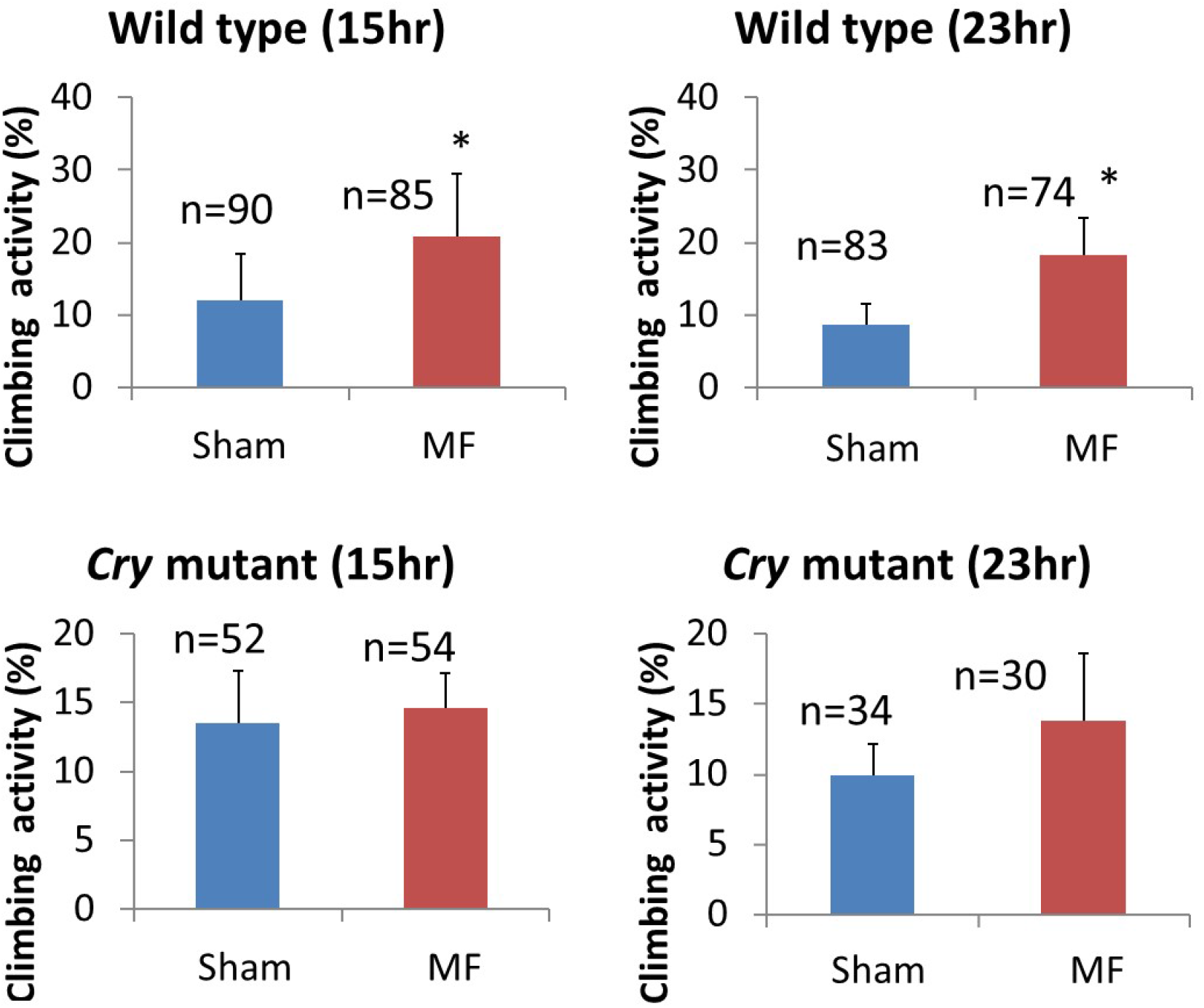
Nocturnal AC magnetic field exposure prevents the decline of motor function. a. Experimental scheme for climbing assay after 3 repeats of AC magnet field exposure during 3 nights. b. Climbing ability of magnetic field (MF)-treated wild type flies was higher than sham-treated ones. But there was no significant difference in *Cry* mutant flies between MF-treated flies and sham-treated flies.

Both wild type and *cry* mutant flies revealed decline in motor function, but surprisingly, AC magnetic field treatment prevented motor function decline in wild type fly. In contrast, AC magnetic field treatment did not exert such effects in *cry* mutant flies (Figure 4b).

## Discussion

Here, using the fruit fly as a magnet receptive model organism, we show that exposure to a specific AC magnetic field during night time affects the health of the fly. *Drosophila melanogaster* is the best model animal, because many mutants and genetically engineered strains are available in world stock centers and laboratories. Here we show that AC magnetic field exposure affected lifespan extension under starvation, sleep improvement and prevention of decreased motor function. In contrast, all the health improvement effect was not observed in *cryptochrome* mutant (*cry*^*b*^) flies suggests that CRY have a key role for the magnet reception in *Drosophila*. Sleep is more than just refreshing, because it is critical for normal body health and function. Interfering with sleep results in problems with lifespan and motor function. In our previous paper, Gaucher’s disease model *Drosophila* showed abnormal motor function, sleep fragmentation and short lifespan^14^. Here we showed that AC magnetic field exposure prevented motor dysfunction in Gaucher’s disease model *Drosophila*.

The data suggest that magnetic field-dependent improvement of sleep quality, lifespan and motor function is mediated by a *cry*-dependent pathway in animals. The *cry* gene has been reported to play a critical role in magnetic detection among diverse species. Genetic studies have shown that a light-dependent magnetic sense in *Drosophila* is mediated by its CRY ^4,21,22^. A recent paper showed using a transgenic approach, that human CRY can function as a magnetosensor in the magnetoreception of *Drosophila* ^4^. Considering these previous data, we thought that all the health improvement molecular mechanisms including lifespan extension, sleep improvement and the prevention of decreased motor function to magnetic field exposure may be involved in the cryptochrome pathway.

## References

1 Vacha, M., Puzova, T. & Kvicalova, M. Radio frequency magnetic fields disrupt magnetoreception in American cockroach. The Journal of experimental biology 212, 3473–3477, doi:10.1242/jeb.028670 (2009).

2 Qin, S. et al. A magnetic protein biocompass. Nature materials 15, 217–226, doi:10.1038/nmat4484 (2016).

3 Fedele, G. et al. Genetic analysis of circadian responses to low frequency electromagnetic fields in Drosophila melanogaster. PLoS genetics 10, e1004804, doi:10.1371/journal.pgen.1004804 (2014).

4 Foley, L. E., Gegear, R. J. & Reppert, S. M. Human cryptochrome exhibits light-dependent magnetosensitivity. Nature communications 2, 356, doi:10.1038/ncomms1364 (2011).

5 Guerra, P. A., Gegear, R. J. & Reppert, S. M. A magnetic compass aids monarch butterfly migration. Nature communications 5, 4164, doi:10.1038/ncomms5164 (2014).

6 Thalau, P., Ritz, T., Stapput, K., Wiltschko, R. & Wiltschko, W. Magnetic compass orientation of migratory birds in the presence of a 1.315 MHz oscillating field. Die Naturwissenschaften 92, 86–90, doi:10.1007/s00114-004-0595-8 (2005).

7 Ritz, T. et al. Magnetic compass of birds is based on a molecule with optimal directional sensitivity. Biophysical journal 96, 3451–3457, doi:10.1016/j.bpj.2008.11.072 (2009).

8 Wang, C. X. et al. Transduction of the Geomagnetic Field as Evidenced from alpha-Band Activity in the Human Brain. eNeuro 6, doi:10.1523/ENEURO.0483-18.2019 (2019).

9 Giachello, C. N., Scrutton, N. S., Jones, A. R. & Baines, R. A. Magnetic Fields Modulate Blue-Light-Dependent Regulation of Neuronal Firing by Cryptochrome. The Journal of neuroscience : the official journal of the Society for Neuroscience 36, 10742–10749, doi:10.1523/JNEUROSCI.2140-16.2016 (2016).

10 Oh, I. T. et al. Behavioral evidence for geomagnetic imprinting and transgenerational inheritance in fruit flies. Proceedings of the National Academy of Sciences of the United States of America 117, 1216–1222, doi:10.1073/pnas.1914106117 (2020).

11 Okano, H. & Ohkubo, C. Effects of neck exposure to 5.5 mT static magnetic field on pharmacologically modulated blood pressure in conscious rabbits. Bioelectromagnetics 26, 469–480, doi:10.1002/bem.20115 (2005).

12 Okano, H., Masuda, H. & Ohkubo, C. Effects of 25 mT static magnetic field on blood pressure in reserpine-induced hypotensive Wistar-Kyoto rats. Bioelectromagnetics 26, 36–48, doi:10.1002/bem.20052 (2005).

13 Xu, S., Okano, H., Tomita, N. & Ikada, Y. Recovery Effects of a 180 mT Static Magnetic Field on Bone Mineral Density of Osteoporotic Lumbar Vertebrae in Ovariectomized Rats. Evidence-based complementary and alternative medicine : eCAM 2011, doi:10.1155/2011/620984 (2011).

14 Kawasaki, H. et al. Minos-insertion mutant of the Drosophila GBA gene homologue showed abnormal phenotypes of climbing ability, sleep and life span with accumulation of hydroxy-glucocerebroside. Gene 614, 49–55, doi:10.1016/j.gene.2017.03.004 (2017).

15 Suzuki, T. et al. Expression of human Gaucher disease gene GBA generates neurodevelopmental defects and ER stress in Drosophila eye. PloS one 8, e69147, doi:10.1371/journal.pone.0069147 (2013).

16 Okano, H., Fujimura, A., Ishiwatari, H. & Watanuki, K. in 2017 IEEE International Conference on Systems, Man, and Cybernetics (SMC). 2442–2447.

17 Kondo, T., Okano, H., Ishiwatari, H. & Watanuki, K. in Advances in Human Factors and Ergonomics in Healthcare and Medical Devices. (ed Nancy J. Lightner) 68–79 (Springer International Publishing).

18 Shaw, P. J., Cirelli, C., Greenspan, R. J. & Tononi, G. Correlates of sleep and waking in Drosophila melanogaster. Science 287, 1834–1837, doi:10.1126/science.287.5459.1834 (2000).

19 Mohite, G. M. et al. Parkinson’s Disease Associated alpha-Synuclein Familial Mutants Promote Dopaminergic Neuronal Death in Drosophila melanogaster. ACS chemical neuroscience 9, 2628–2638, doi:10.1021/acschemneuro.8b00107 (2018).

20 Feany, M. B. & Bender, W. W. A Drosophila model of Parkinson’s disease. Nature 404, 394–398, doi:10.1038/35006074 (2000).

21 Gegear, R. J., Casselman, A., Waddell, S. & Reppert, S. M. Cryptochrome mediates light-dependent magnetosensitivity in Drosophila. Nature 454, 1014–1018, doi:10.1038/nature07183 (2008).

22 Yoshii, T., Ahmad, M. & Helfrich-Forster, C. Cryptochrome mediates light-dependent magnetosensitivity of Drosophila’s circadian clock. PLoS biology 7, e1000086, doi:10.1371/journal.pbio.1000086 (2009).

